# Typical lipreading and audiovisual speech perception without motor simulation

**DOI:** 10.1101/2020.06.03.131813

**Authors:** Gilles Vannuscorps, Michael Andres, Sarah Carneiro, Elise Rombaux, Alfonso Caramazza

## Abstract

All it takes is a face to face conversation in a noisy environment to realize that viewing a speaker’s lip movements contributes to speech comprehension. Following the finding that brain areas that control speech production are also recruited during lip reading, the received explanation is that lipreading operates through a covert unconscious imitation of the observed speech movements in the observer’s own speech motor system – a motor simulation. However, motor effects during lipreading do not necessarily imply simulation or a causal role in perception. In line with this alternative, we report here that some individuals born with lip paralysis, who are therefore unable to covertly imitate observed lip movements, have typical lipreading abilities and audiovisual speech perception. This constitutes existence proof that typically efficient lipreading abilities can be achieved without motor simulation. Although it remains an open question whether this conclusion generalizes to typically developed participants, these findings demonstrate that alternatives to motor simulation theories are plausible and invite the conclusion that lip-reading does not involve motor simulation. Beyond its theoretical significance in the field of speech perception, this finding also calls for a re-examination of the more general hypothesis that motor simulation underlies action perception and interpretation developed in the frameworks of the motor simulation and mirror neuron hypotheses.

## Introduction

In face to face conversations, speech perception results from the integration of the outcomes of an auditory analysis of the sounds that the speaker produces and of the visual analysis of her lip and mouth movements (McDonald & McGurk, 1978; McGurk & McDonald, 1976; Reisberg, McLean & Goldfield, 1987; Sumby & Pollack, 1954). When auditory and visual information are experimentally made incongruent (e.g., an auditory bilabial /ba/ and a visual velar /ga/), for instance, participants often report perceiving an intermediate consonant (e.g., in this case the dental /da/) (McDonald & McGurk, 1978; McGurk & McDonald, 1976; Massaro, 1989).

Following the observation that participants asked to silently lipread recruit not only parts of their visual system but also the inferior frontal gyrus and the premotor cortex typically involved during the execution of the same facial movements (Callan et al., 2003; Calvert & Campbell, 2003; Okada & Hickok, 2009; Sato, Buccino, Gentilucci & Cattaneo, 2010; Skipper, Nusbaum, & Small, 2005; Watkins, Strafella & Paus, 2003), the received explanation has been that the interpretation of visual speech requires a covert unconscious imitation of the observed speech movements in the observer’s motor system – a motor simulation of the observed speech gestures. This internal “simulated” speech production would allow interpreting the observed movements and, therefore, constrains the interpretation of the acoustic signal (Callan et al., 2003; Callan et al., 2004; Okada & Hickok, 2009; Chu et al., 2013; Skipper, Goldin-Meadows, Nusbaum & Small, 2007; Skipper, Van Wassenhove, Nusbaum & Small, 2007; Tye-Murray, Spehar, Myerson, Hale & Sommers, 2013).

However, these findings are open to alternative interpretations. For instance, the activation of the motor system could be a consequence, rather than a cause, of the identification of the visual syllables. This interpretation implies that the visual analysis of lip movements provides sufficient information to accurately lipread and does not require the contribution of the motor system. To help choose between these hypotheses, we compared word-level lipreading abilities (in Experiment 1) and the strength of the influence of visual speech on the interpretation of auditory speech information (in Experiment 2) in typically developed participants and in eleven individuals born with congenitally reduced or completely absent lip movements in the context of the Moebius syndrome (individuals with the Moebius syndrome, IMS, see table 1) - an extremely rare congenital disorder characterized, among other things, by a non-progressive facial paralysis caused by an altered development of the facial (VII) cranial nerve (Verzijl, Van der Zwaag, Cruysberg & Padberg, 2003).

**Table 1.** Demographic and clinical data of the individuals with the Moebius Syndrome

|  | Sex | Age<br>(years) | Education<br>(years) | Mid-level<br>perception<sup>1</sup> | Inferior lip | Superior lip |
| --- | --- | --- | --- | --- | --- | --- |
| IMS1 | F | 37 | 4 | 0.9 | No movements | No movements |
| IMS2 | F | 36 | 3 | -3.3 | Slight movements | No movements |
| IMS3 | M | 19 | 2 | -0.3 | Slight movements | No movements |
| IMS4 | F | 43 | 1 | -2.9 | Slight movements | No movements |
| IMS5 | F | 31 | 2 | -0.3 | Slight movements | Slight movements |
| IMS6 | F | 19 | 0 | 0.1 | Mild movements | No movements |
| IMS7 | M | 31 | 2 | 0.5 | No movements | No movements |
| IMS8 | F | 21 | 3 | 0.5 | Slight movements | No movements |
| IMS9 | F | 15 | 0 | -3.3 | Mild movements | No movements |
| IMS10 | F | 20 | 0 | -0.3 | No movements | No movements |
| IMS11 | M | 33 | 3 | -4.2 | No movements | No movements |

<sup>1</sup> Participants' modified *t*-test (Crawford & Howell, 1998) at the Leuven Perceptual Organization Screening Test, L-POST (Torfs, Vancleef, Lafosse, Wagemans & de Wit, 2014).

Three main arguments support the assumption that a congenital lip paralysis prevents simulating observed lip movements to help lip-reading. First, extant evidence suggests that the motor cortex does not contain representations of congenitally absent or deeferented limbs (e.g., Reilly & Sirigu, 2011). Rather, the specific parts of the somatosensory and motor cortices that would normally represent the “absent” or deeferented limbs are allocated to the representation of adjacent body parts (Funk et al., 2008; Kaas, 2000; Kaas, Merzenich & Killackey, 1983, 1983; Makin, Scholtz, Henderson Slater, Johansen-Berg, & Tracey, 2015; Stoeckel, Seitz, & Buethefish, 2009; Striem-Amit, Vannuscorps & Caramazza, 2017). Beyond this evidence, in any event, it is unclear how a motor representation of lip movement could be formed in individuals who have never had the ability to execute any lip movement, and we are not aware of any attempt at describing how such a mechanism would operate. Furthermore, it is unclear in what sense such a representation would be a “motor representation”. Second, merely “having” lip motor representations would not be sufficient to simulate an observed lip movement. Motor simulation is not based merely on motor representations of body parts (e.g., of the lips) but on representations of the movements previously executed with these body parts. Hence, previous motor experience with observed body movements is critical for motor simulation to occur (e.g., Calvo-Merino, Grezès, Glaser, Passingham & Haggard, 2006; Swaminathan et al., 2013; Turella, Wurm, Tucciarelli & Lingnau, 2013) and lipreading efficiency is assumed to depend on the similarity between the observed lip movements and those used by the viewer (Tye-Murray, Spehar, Myerson, Hale & Sommers, 2013; 2015). Since the IMS have never executed lip movements, it is unclear how they could motorically simulate observed lip movements, such as a movement of the lower lip to contact the upper teeth rapidly followed by an anteriorization, opening and rounding of the two lips involved in the articulation of the syllable (/fa/). Third, in any event, a motor simulation of observed lip movements by the IMS would not be sufficient to support the IMS’s lip reading abilities according to motor simulation theories. According to these theories, motor simulation of observed body movements is necessary but not sufficient to support lipreading. The role of motor simulation derives from the fact that it is supposed to help retrieving information about the observed movements acquired through previous motor experience. Motor simulation, for instance, supports lipreading because simulating a given observed lip movement allows the observer to “retrieve” what sound or syllable these movements allow him/her to produce when s/he carries out that particular motor program. Since the IMS have never themselves generated the lip movements probed in this study, motor simulation could not be regarded as a possible support to lipreading.

In the light of these considerations, the investigation of the contribution of lip reading on the perception of speech in IMS provides a clear test of the motor simulation hypothesis: if this hypothesis is valid, then, it should be impossible for any individual deprived of lip motor representations to be as efficient as controls at lip reading.

It is important to note, however, that Moebius syndrome typically impacts not only the individuals’ sensorimotor system, but also their visual, perceptual, cognitive, and social abilities to various extents (Bate et al., 2013; Carta, Mora, Neri, Favilla, & Sadun, 2011; Vannuscorps, Andres & Caramazza, 2020). This has significant interpretative and methodological implications for the current study. Given the complexity of the disorder, it is not unexpected that at least some IMS would show relatively poor lipreading performance. Determining the cause of the lipreading deficit in these individuals is not a straightforward matter: candidate causes include not only their production disorder but other impaired but functionally separate processes, such as visuo-perceptual processing, which co-occur to varying degrees in these individuals. This situation creates an asymmetry in the evidentiary value of good versus poor lipreading performance: normotypical performance on this task indicates that the absence of motor simulation is not necessary for lipreading, whereas poor performance is indeterminate on the role of motor simulation in lipreading. This interpretative asymmetry has a methodological consequence in the context of the specific prediction tested in this study – that none of the individuals with congenital facial paralysis should be as efficient as the controls in lipreading. The appropriate methodology is the use of single-case analyses, since this approach allows us to determine unambiguously whether the inability to carry out the relevant motor simulation necessarily adversely affects lipreading performance.

## Methods

The experimental investigations were carried out from October 2015 to September 2019 in sessions lasting between 60 and 90 minutes. The study was approved by the biomedical ethics committee of the Cliniques Universitaires Saint-Luc, Brussels, Belgium, was performed in accordance with relevant guidelines/regulations and all participants gave written informed consent prior to the study and the research.

The experiments were controlled by the online testable.org interface (http://www.testable.org), which allows precise spatiotemporal control of online experiments. Control participants were tested on the 15.6-inch anti-glare screen (set at 1366 x 768 pixels and 60Hz) of a Dell Latitude E5530 laptop operated by Windows 10. The individuals with the Moebius Syndrome (IMS) were tested remotely on their own computer under supervision of the experimenter through a visual conference system. At the beginning of each experiment, the participant was instructed to set the browsing window of the computer to full screen, minimize possible distractions (e.g., TV, phone, etc.) and position themselves at arm’s length from the monitor for the duration of the experiment. Next, a calibration procedure ascertaining homogeneous presentation size and time on all computer screens took place. Next, participants started the experiment.

### Participants

Eleven individuals with congenitally reduced or completely absent lip movements in the context of the Moebius Syndrome (individuals with the Moebius Syndrome, IMS, see table 1 and Vannuscorps, Andres & Caramazza, 2020) and 25 typically developed highly educated young adults (15 females; 3 left-handed; all college students or graduates without any history of psychiatric or neurological disorder; Mean age ± SD: 28.6 ± 6.5 years) participated in Study 1. Eight of the IMS (IMS1, 3, 4, 5, 8, 9, 10, 11) and 20 new typically developed highly educated young adults (13 females; 2 left-handed; all college students or graduates without any history of psychiatric or neurological disorder; Mean age ± SD: 22.7 ± 2.3 years) participated in Study 2.

The participants with the Moebius Syndrome included in this study presented with congenital bilateral facial paralyses of different degrees of severity. As indicated in Table 1, lip movements were completely absent in IMS 1, 7, 10 and 11, very severely reduced in IMS 2, 3 and 4, and severely reduced in IMS 5, 6, 8 and 9. IMS 2 could only very slightly pull her lower lip downward. IMS 3 could very slightly pull the corners of his mouth down. This movement was systematically accompanied by a slight increase of mouth opening and a wrinkling of the surface of the skin of the neck, suggesting a slight contraction of the platysma muscle. IMS 4 could slightly contract her cheeks (buccinators), resulting in a slight upward and stretching of the lips. In all these participants, there was no mouth closing, no movement of the superior lip and none of these individuals were able to move their lips in a position corresponding to bilabial (/p/, /b/, /m/) or labiodental (/f/, /v/) French consonants or to rounded 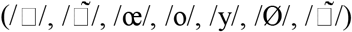 and stretched (/i/, /e/) French vowels. IMS 5, 6, 8 and 9 were able to open and close the mouth. IMS 5 could also slightly pull the angles of the mouth backwards by contracting the cheeks, slightly pull her lower lip downwards and very slightly contract the right sight of her upper lip. IMS 6 could also pull the angles of the mouth backwards by contracting the cheeks. IMS 8 was able to execute a mild combined backward/upward movement of the right angle of the mouth and a slight backward movement of the left angle of the mouth. IMS9 could normally pull the right angles of the mouth backwards by contracting the right cheek and slightly pull the left angles of the mouth backwards by contracting the left cheek. IMS 5, 6, 8 and 9 were thus able to move their lips in a position corresponding to bilabial French consonants and to stretched French vowels but not in a position corresponding to labiodental consonants and rounded vowels.

It is important to note that in addition to these motor symptoms, the individuals with the Moebius syndrome typically also present with a heterogeneous spectrum of visuo-perceptual disorders, including various patterns of ocular motility alterations, various degrees of visual acuity impairments, of lagophthalmos, an absence of stereopsis, and frequent mid and high visual perception problems (Verzijl, Van der Zwaag, Cruysberg & Padberg, 2003; Bate, Cook, Mole & Cole, 2013; Calder, Keane, Cole, Campbell & Young, 2000). The performance of the IMS participants included in this study in a mid-level perception screening test ranged from typical to severely impaired (see Table 1), for example.

### Stimuli and procedure

#### Experiment 1

Stimuli were 60 silent video-clips (~3 seconds, 30 frames/second, 854 x 480 pixels) showing one of two actresses articulating a word in French (see list in appendix). Only the face of the actress was visible. Each video started with the actress in a neutral posture, followed by the articulation and ended by a return in the neutral posture. Each video was associated with a target word and either 2 (for consonants, N = 37) or 3 (for vowels, N = 23) distractors. For 23 of the video-clips, 3 distractors differing from the target word in terms of a single vowel articulated with a different shape and/or aperture of the mouth were selected. For instance, the words “plan” (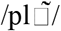, i.e., an open, rounded vowel), “plot” (/plo/, i.e., a mid-open, rounded vowel) and “pli” (/pli/, i.e., a closed, non-rounded vowel) were used as response alternatives for the target word “plat” (/pla/, i.e., an open, non-rounded vowel). For the remaining 37 video-clips, 2 distractors differing from the target word in terms of the place of articulation of a single consonant were selected. For instance, the words “fente” (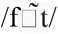, i.e., a labiodental consonant) and “chante” (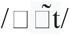, i.e., a postalveolar consonant) were selected as response alternatives for the target word “menthe” (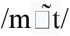, i.e., a bilabial consonant).

Each of the 60 trials started with the presentation of three or four words on the computer screen for 5 sec, followed by the presentation of a video-clip of an actress articulating one of the words lasting ± 3 seconds and three or four response buttons. Participants were asked to read the words carefully, observe the video carefully and, then, to identify the word articulated by the actress by clicking on the corresponding response button. There was no time constraint for responding but participants were asked not to respond before the end of the video clip.

#### Experiment 2

Stimuli were 32 video-clips (1.5 seconds, 50 frames/second, 960 x 544 pixels) showing one of four actors (two males, two females) articulating twice in a row one of six syllables paired with a congruent audio (« pa », « ta », « ka », « ba », « da » and « ga ») or one of two syllables paired with an incongruent audio (visual « ga » and « ka » paired with the audio /ba/ and /pa/, respectively) and 8 similar video-clips in which a small pink dot appeared at a random place on the face of the actor (one by condition, 2 by actor). Only the shoulders and face of the actors/actresses was visible. Each video started with the actor in a neutral posture, followed by the articulation and ended by a return in the neutral posture. The actors maintained an even intonation, tempo, and vocal intensity while producing the syllables.

During the experiment, participants were first presented with an auditory-only stimulus and asked to set the volume of their computer at a clearly audible, comfortable level. Then, they received the following instruction (translated from French): “In this experiment, we will test your ability to do two tasks simultaneously. You will see video-clips of actors articulating twice the same syllable. On some of these video-clips, a small pink dot will appear somewhere on the actor’s face. After each video, we ask you to click on the response button “pink dot” if you have seen a small pink dot. If no small pink dot has appeared, then simply report the syllable that you heard by clicking on the corresponding syllable on the computer screen.” After the instructions, participants saw, in a pseudo-random order, a series of 128 video-clips comprising 3 repetitions of each actor articulating twice the same syllable paired with a congruent audio (3 repetition x 4 actors x 6 congruent stimuli = 72 video-clips), 6 repetitions of each actor articulating twice the same syllable paired with an incongruent audio (6 repetition x 4 actors x 2 congruent stimuli = 48 video-clips) and 2 videos of each actor articulating twice the same syllable paired with a congruent audio in which a small pink dot appeared on the actors’ face. After each video-clip the participant was asked to indicate if s/he had seen a pink dot and, if not, to click on the syllable articulated by the actor presented among 6 alternatives (“pa”, “ta”, “ka”, “ba”, “da”, “ga”). There was no time constraint for responding.

### Analysis and results

Given the heterogeneity of the clinical expression of Moebius Syndrome, especially in terms of associated visuo-perceptual symptoms (Bate et al., 2013; Carta et al., 2011; see Table 1), and the specific prediction of the motor simulation hypothesis tested in this study – viz., that none of the individuals with congenital facial paralysis should be as efficient as the controls in lip reading – we conducted analyses focused on the performance of each IMS. We used Crawford and Howell’s (1998) modified t-test to compare the performance of each IMS to that of the control group and establish whether it is possible for an individual with congenital lip paralysis to achieve normotypical efficient lip-reading skills. To minimize the likelihood of false negatives, that is, the risk to conclude erroneously that an IMS achieves a “normotypical level” of efficiency in a lipreading experiment, we selected a high alpha level *(p >* 0.2), which set the threshold for “typically efficient” performance in a given experiment to scores above 0.85 standard deviations below the mean of the controls, *after* control participants with an abnormally low score were dismissed.

Experiment 1 tested the hypothesis that motor simulation is necessary for efficient lipreading. We counted the number of correct responses of each participant (see Figure 1A). Unsurprisingly given the visuo-perceptual symptoms commonly associated with Moebius Syndrome, four IMS (2, 3, 9, 11) performed below, or among the less efficient control participants. More interestingly, and contrary to expectations from the motor simulation hypothesis, the other seven IMS performed above the mean performance of the controls despite severely reduced (IMS 5, 6, 8), very severely reduced (IMS 4) or even completely absent (IMS 1, 7, 10) lip motor representations (all modified *t-*tests, *t*s (24) > 0).

**Figure 1.**
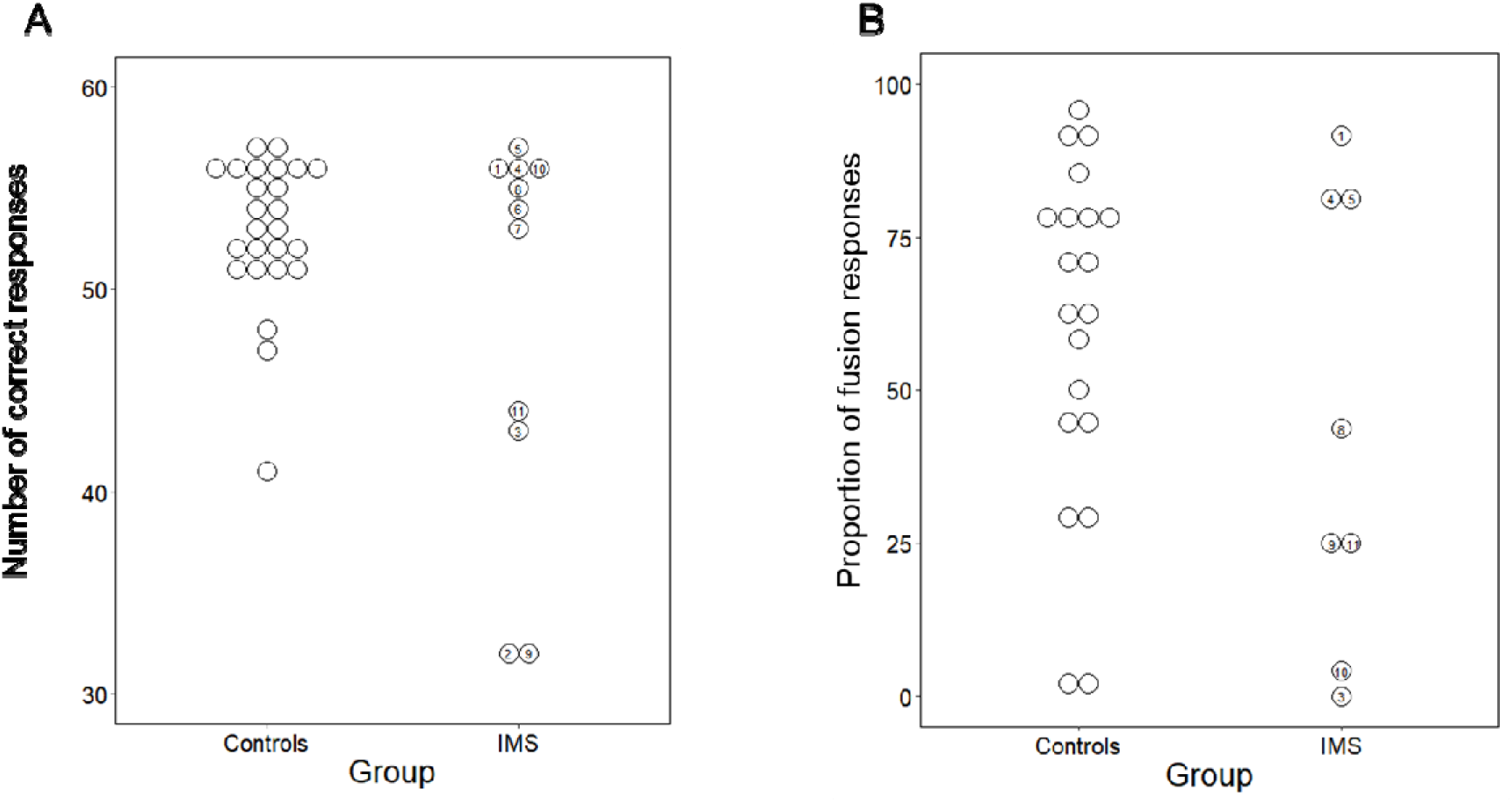
Results of Experiment 1 (A) and 2 (B) by group and by individual participant. The numbers refer to the IMS reported in table 1. The proportion of fusion responses reported in panel B corresponds to the percentage of trials in which incongruent pairings between visual velar and auditory bilabial consonants (N=48) produced the percept of intermediate dental consonants (/da/ and /ta/).

Experiment 2 tested the hypothesis that motor simulation underlies typical audio-visual integration for speech perception. In such experiments, the influence of visual upon auditory speech is indexed by the proportion of trials in which incongruent pairings between visual velar and auditory bilabial consonants produced the percept of intermediate dental consonants (/da/ and /ta/), i.e., “fusion responses” (McDonald & McGurk, 1978; McGurk & McDonald, 1976). Thus, we computed the proportion of fusion responses of each participant (Figure 1B) as an index of the contribution of the visual signal on the perception of the auditory syllables. Contrary to expectations from the motor simulation hypothesis, three IMS reported perceiving a large proportion of fusion responses despite (very) severely reduced (IMS 4, 5) or even completely absent (IMS 1) lip motor representations (all modified *t-*tests, *t*s (19) > 0). Despite completely absent lip motor representations, for instance, IMS1 had 92% of fusion responses, a proportion that was larger for only one of the control participants.

## Discussion

We tested a prediction derived from the hypothesis that efficient lipreading and typical audiovisual speech perception require covert imitation — motor simulation — of the observed speech movements. In two experiments we found that several participants with congenital facial paralysis were as good at lipreading (Experiment 1) and as influenced by visual speech (Experiment 2) as control participants despite severely reduced or even completely absent lip motor representations. IMS 1, for instance, had a congenital complete lip paralysis but was nevertheless better at lipreading than 17/25 control participants in Experiment 1 and perceived a larger proportion of fusion responses than 17/20 control participants in Experiment 2. It follows that (1) typical lipreading efficiency does not require motor simulation and, as a corollary, that (2) the visual system can independently support efficient lip reading.

In both experiments some individuals with a congenital facial paralysis performed significantly below the control participants. The finding that other individuals with an equally or even more severe motor disorder achieved a normal level of performance in these two tasks forces the conclusion that their more marked difficulties in lipreading cannot be explained by their motor disorder. Convergent evidence for this conclusion is provided by the fact that three of the four IMS participants who performed significantly less accurately than the controls in Experiment 1 also performed significantly below control participants in a visual perceptual screening test (see Table 1), suggesting that their difficulties are likely the consequence of a visuo-perceptual deficit.

In line with our conclusion, previous studies had already reported that lipreading abilities precede speech production abilities during development. For instance, two- to five-month-old infants who have not yet mastered articulated speech look longer at a face executing articulatory gestures matching a simultaneously presented sound than at a face that does not (Kuhl & Meltzoff, 1982; Patterson & Werker, 1999; Patterson & werker, 2003). These studies demonstrated that it is possible to develop some level of lipreading capability without motor simulation. Our findings additionally demonstrate that it is possible to reach normal lipreading efficiency without motor simulation.

These findings support the hypothesis that lip reading is a property of the visuo-perceptual system unaided by the motor system (Bernstein & Liebenthal, 2014; Matchin, Groulx & Hickok, 2014). According to this view, lipreading requires a visuo-perceptual analysis of the actor’s configural and dynamic facial features to provide access to stored visual descriptions of the facial postures and movements corresponding to different linguistic units. Once this stored visual representation is accessed, it may be integrated with auditory information in multisensory integration sites to support audiovisual speech comprehension (Beauchamp, Argall, Borduka, Duyn & Martin, 2004).

Our results demonstrate that it is possible to account for typically efficient lipreading abilities without appealing to motor simulation. Admittedly, it is possible that lipreading in the IMS relies on atypical mechanisms and, therefore, it is an open question whether our conclusion generalizes to typically developed participants. Future studies are needed to elucidate this question with the help of neuropsychological studies of patients suffering from brain damage that affects their ability to imitate lip movements covertly. Nevertheless, there seems to be currently no compelling empirical reason to favor the less parsimonious motor simulation hypothesis. Hence, our findings at the very least emphasizes the need for a shift in the burden of proof relative to the question of the role of motor simulation in lipreading.

This conclusion converges with that of previous reports of IMS participants who achieved normal levels of performance in facial expression recognition despite their congenital facial paralysis (Bate et al., 2013; Calder et al., 2000; Bogart & Matsumoto, 2010) and of individuals congenitally deprived of hand motor representations who nonetheless perceived and comprehended hand actions as efficiently and with the same biases and brain networks as typically developed participants (Vannuscorps, Pillon & Andres, 2012; Vannuscorps, Andres & Pillon, 2013; Vannuscorps & Caramazza, 2015, 2016a, b, 2017, 2019; Vannuscorps, Wurm, Striem-Amit & Caramazza, 2019). Together, these results severely challenge the hypothesis that body movement perception and comprehension rely on motor simulation (Rizzolatti & Sinigaglia, 2010).

## Acknowledgments

This research was supported by the Mind, Brain and Behavior Interfaculty Initiative provostial funds. MA is a research associate at the Fonds National de la Recherche Scientifique (FRS-FNRS, Belgium). We would like to thank the Association Syndrome Moebius France for its help all along this project.

## Appendix 1

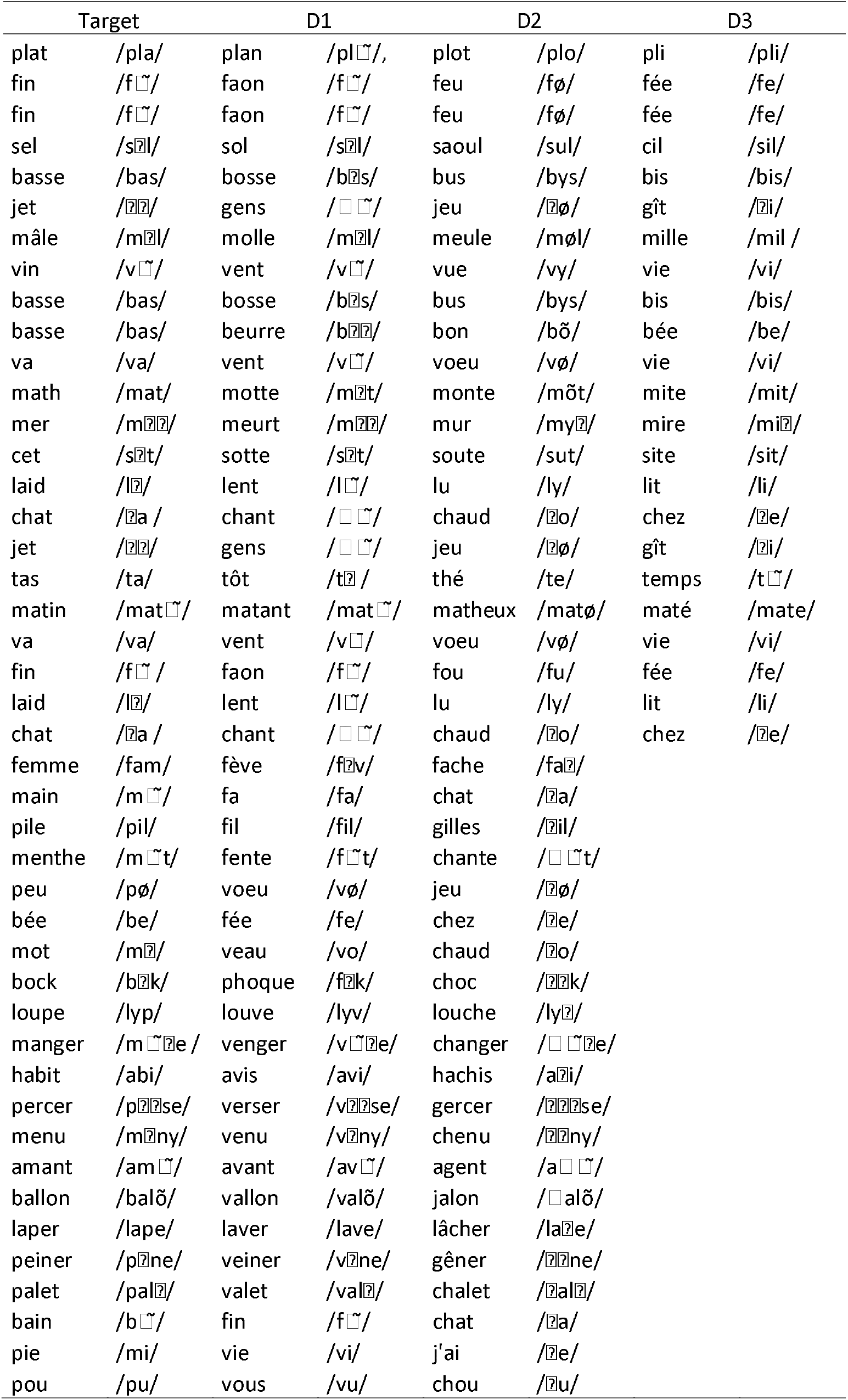
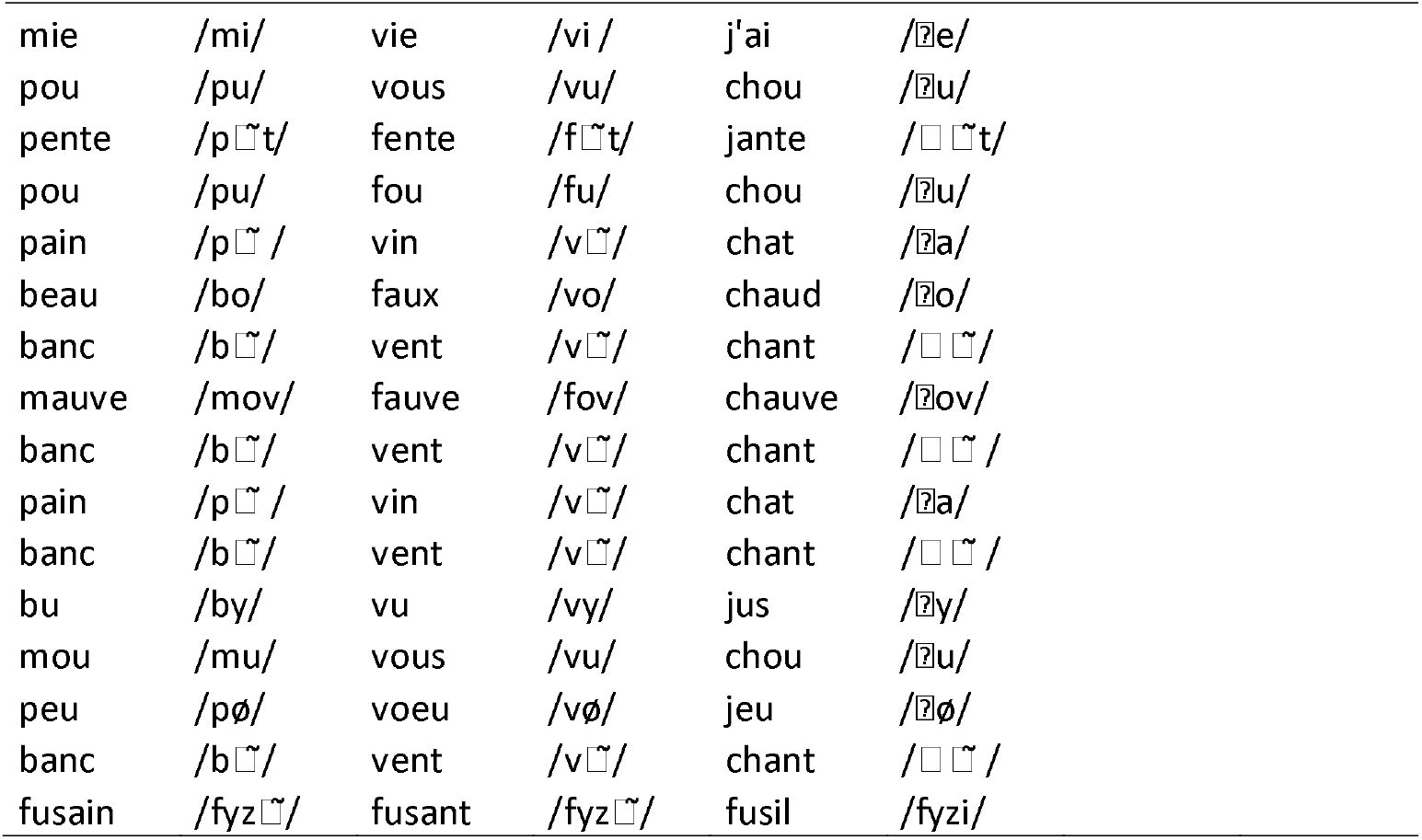
Orthographic and phonetic transcription of the target words articulated by the actresses in Experiment 1 and the associated distractor stimuli.

## References

Bate, S., Cook, S. J., Mole, J., & Cole, J. (2013). First Report of Generalized Face Processing Difficulties in Möbius Sequence. PLoS ONE, 5(4). https://doi.org/10.1371/journal.pone.0062656

Beauchamp, M. S., Argall, B. D., Bodurka, J., Duyn, J. H., & Martin, A. (2004). Unraveling multisensory integration: Patchy organization within human STS multisensory cortex. Nature Neuroscience, 7(11), 1190–1192. https://doi.org/10.1038/nn1333

Bernstein, L. E., & Liebenthal, E. (2014). Neural pathways for visual speech perception. Frontiers in Neuroscience. Frontiers Media S.A. https://doi.org/10.3389/fnins.2014.00386

Bogart, K. R., & Matsumoto, D. (2010). Facial mimicry is not necessary to recognize emotion: Facial expression recognition by people with Moebius syndrome. Social Neuroscience, 5(2), 241–251. https://doi.org/10.1080/17470910903395692

Calder, A. J., Keane, J., Cole, J., Campbell, R., & Young, A. W. (2000). Facial expression recognition by people with mobius syndrome. Cognitive Neuropsychology, 17(1-3), 73–87. https://doi.org/10.1080/026432900380490

Callan, D. E., Jones, J. A., Munhall, K., Callan, A. M., Callan, A. M., & Vatikiotis-Bateson, E. (2003). Neural processes underlying perceptual enhancement by visual speech gestures. NeuroReport, 14(17), 746–748. https://doi.org/10.1097/00001756-200312020-00016

Callan, D. E., Jones, J. A., Munhall, K., Kroos, C., Callan, A. M., & Vatikiotis-Bateson, E. (2004). Multisensory integration sites identified by perception of spatial wavelet filtered visual speech gesture information. Journal of Cognitive Neuroscience, 16(5), 805–816. https://doi.org/10.1162/089892904970771

Calvert, G. A., & Campbell, R. (2003). Reading speech from still and moving faces: The neural substrates of visible speech. Journal of Cognitive Neuroscience, 15(1), 57–70. https://doi.org/10.1162/089892903321107828

Calvo-Merino, B., Grèzes, J., Glaser, D. E., Passingham, R. E., & Haggard, P. (2006). Seeing or Doing? Influence of Visual and Motor Familiarity in Action Observation. Current Biology, 16(19), 1905–1910. https://doi.org/10.1016/j.cub.2006.07.065

Carta, A., Mora, P., Neri, A., Favilla, S., & Sadun, A. A. (2011). Ophthalmologic and systemic features in Möbius syndrome: An Italian case series. Ophthalmology, 118(8), 1518–1523. https://doi.org/10.1016/j.ophtha.2011.01.023

Chu, Y. H., Lin, F. H., Chou, Y. J., Tsai, K. W. K., Kuo, W. J., & Jääskeläinen, I. P. (2013). Effective cerebral connectivity during silent speech reading revealed by functional magnetic resonance imaging. PLoS ONE, 8(11). https://doi.org/10.1371/journal.pone.0080265

Crawford, J. R., & Howell, D. C. (1998). Comparing an individual’s test score against norms derived from small samples. Clinical Neuropsychologist, 12(4), 482–486. https://doi.org/10.1076/clin.12.4.482.7241

Funk, M., Lutz, K., Hotz-Boendermaker, S., Roos, M., Summers, P., Brugger, P., … Kollias, S. S. (2008). Sensorimotor tongue representation in individuals with unilateral upper limb amelia. NeuroImage, 43(1), 121–127. https://doi.org/10.1016/j.neuroimage.2008.06.011

Kaas, J. H. (2000). The reorganization of somatosensory and motor cortex after peripheral nerve or spinal cord injury in primates. Progress in Brain Research, 128, 173–179.

Kaas, J. H., Merzenich, M. M., & Killackey, H. P. (1983). The Reorganization of Somatosensory Cortex Following Peripheral Nerve Damage in Adult and Developing Mammals. Annual Review of Neuroscience, 6(1), 325–356. https://doi.org/10.1146/annurev.ne.06.030183.001545

Kuhl, P. K., & Meltzoff, A. N. (1982). The bimodal perception of speech in infancy. Science, 218(4577), 1138–1141. https://doi.org/10.1126/science.7146899

MacDonald, J., & McGurk, H. (1978). Visual influences on speech perception processes. Perception & Psychophysics, 24(3), 253–257. https://doi.org/10.3758/BF03206096

Makin, T. R., Scholz, J., Henderson Slater, D., Johansen-Berg, H., & Tracey, I. (2015). Reassessing cortical reorganization in the primary sensorimotor cortex following arm amputation. Brain, 138(8), 2140–2146. https://doi.org/10.1093/brain/awv161

Massaro, D. W. (1989). Multiple Book Review of Speech perception by ear and eye: A paradigm for psychological inquiry. Behavioral and Brain Sciences, 12(4), 741–755. https://doi.org/10.1017/s0140525x00025619

Matchin, W., Groulx, K., & Hickok, G. (2014). Audiovisual speech integration does not rely on the motor system: Evidence from articulatory suppression, the McGurk effect, and fMRI. Journal of Cognitive Neuroscience, 26(3), 606–620. https://doi.org/10.1162/jocn_a_00515

McGurk, H., & Macdonald, J. (1976). Hearing lips and seeing voices. Nature, 264(5588), 746–748. https://doi.org/10.1038/264746a0

Okada, K., & Hickok, G. (2009). Two cortical mechanisms support the integration of visual and auditory speech: A hypothesis and preliminary data. Neuroscience Letters, 452(3), 219–223. https://doi.org/10.1016/j.neulet.2009.01.060

Patterson, M. L., & Werker, J. F. (1999). Matching phonetic information in lips and voice is robust in 4.5-month-old infants. Infant Behavior and Development, 22(2), 237–247. https://doi.org/10.1016/S0163-6383(99)00003-X

Patterson, M. L., & Werker, J. F. (2003). Two-month-old infants match phonetic information in lips and voice. Developmental Science, 6(2), 191–196. https://doi.org/10.1111/1467-7687.00271

Reisberg, D., McLean, J., & Goldfield, A. (1987). Easy to hear but hard to understand: A lipreading advantage with intact auditory stimuli. In B. Dodd & R. Campbell (Eds.), Hearing by Eye: The Psychology of Lip-reading (pp. 97–113). Hillsdale, NJ: Erlbaum.

Reilly, K. T., & Sirigu, A. (2011). Motor cortex representation of the upper-limb in individuals born without a hand. PLoS ONE, 6(4). https://doi.org/10.1371/journal.pone.0018100

Rizzolatti, G., & Sinigaglia, C. (2010). The functional role of the parieto-frontal mirror circuit: Interpretations and misinterpretations. Nature Reviews Neuroscience. https://doi.org/10.1038/nrn2805

Sato, M., Buccino, G., Gentilucci, M., & Cattaneo, L. (2010). On the tip of the tongue: Modulation of the primary motor cortex during audiovisual speech perception. Speech Communication, 52(6), 533–541. https://doi.org/10.1016/j.specom.2009.12.004

Skipper, J. I., Goldin-Meadow, S., Nusbaum, H. C., & Small, S. L. (2007). Speech-associated gestures, Broca’s area, and the human mirror system. Brain and Language, 101(3), 260–277. https://doi.org/10.1016/j.bandl.2007.02.008

Skipper, J. I., Van Wassenhove, V., Nusbaum, H. C., & Small, S. L. (2007). Hearing lips and seeing voices: How cortical areas supporting speech production mediate audiovisual speech perception. Cerebral Cortex, 17(10), 2387–2399. https://doi.org/10.1093/cercor/bhl147

Skipper, J. I., Nusbaum, H. C., & Small, S. L. (2005). Listening to talking faces: Motor cortical activation during speech perception. NeuroImage, 25(1), 76–89. https://doi.org/10.1016/j.neuroimage.2004.11.006

Stoeckel, M. C., Seitz, R. J., & Buetefisch, C. M. (2009). Congenitally altered motor experience alters somatotopic organization of human primary motor cortex. Proceedings of the National Academy of Sciences of the United States of America, 106(7), 2395–2400. https://doi.org/10.1073/pnas.0803733106

Striem-Amit, E., Vannuscorps, G., & Caramazza, A. (2018). Plasticity based on compensatory effector use in the association but not primary sensorimotor cortex of people born without hands. Proceedings of the National Academy of Sciences of the United States of America, 115(30), 7801–7806. https://doi.org/10.1073/pnas.1803926115

Sumby, W. H., & Pollack, I. (1954). Visual Contribution to Speech Intelligibility in Noise. Journal of the Acoustical Society of America, 26(2), 212–215. https://doi.ore/10.1121Z1.1907309

Swaminathan, S., MacSweeney, M., Boyles, R., Waters, D., Watkins, K. E., & Möttönen, R. (2013). Motor excitability during visual perception of known and unknown spoken languages. Brain and Language, 126(1), 1–7. https://doi.org/10.1016/j.bandl.2013.03.002

Torfs, K., Vancleef, K., Lafosse, C., Wagemans, J., & de-Wit, L. (2014). The Leuven Perceptual Organization Screening Test (L-POST), an online test to assess mid-level visual perception. Behavior Research Methods, 46(2), 472–487. https://doi.org/10.3758/s13428-013-0382-6

Turella, L., Wurm, M. F., Tucciarelli, R., & Lingnau, A. (2013). Expertise in action observation: Recent neuroimaging findings and future perspectives. Frontiers in Human Neuroscience, (OCT). https://doi.org/10.3389/fhhum.2013.00637

Tye-Murray, N., Spehar, B. P., Myerson, J., Hale, S., & Sommers, M. S. (2013). Reading your own lips: Common-coding theory and visual speech perception. Psychonomic Bulletin and Review, 20(1), 115–119. https://doi.org/10.3758/s13423-012-0328-5

Tye-Murray, N., Spehar, B. P., Myerson, J., Hale, S., & Sommers, M. S. (2015). The self-advantage in visual speech processing enhances audiovisual speech recognition in noise. Psychonomic Bulletin and Review, 22(4), 1048–1053. https://doi.org/10.3758/s13423-014-0774-3

Vannuscorps, G., & Caramazza, A. (2015). Typical biomechanical bias in the perception of congenitally absent hands. Cortex, 67(147), e150.

Vannuscorps, G., & Caramazza, A. (2016a). The origin of the biomechanical bias in apparent body movement perception. Neuropsychologia, 89, 281–286. https://doi.org/10.1016/j.neuropsychologia.2016.05.029

Vannuscorps, G., & Caramazza, A. (2016b). Typical action perception and interpretation without motor simulation. Proceedings of the National Academy of Sciences of the United States of America, 113(1), 86–91. https://doi.org/10.1073/pnas.1516978112

Vannuscorps, G., & Caramazza, A. (2017). Typical predictive eye movements during action observation without effector-specific motor simulation. Psychonomic Bulletin & Review, 24(4), 1152–1157.

Vannuscorps, G., & Caramazza, A. (2019). Conceptual processing of action verbs with and without motor representations. Cognitive Neuropsychology, 36(7-8), 301–312.

Vannuscorps, G., Andres, M., & Caramazza, A. (2020). Efficient recognition of facial expressions does not require motor simulation. Elife, 9, e54687.

Vannuscorps, G., Andres, M., & Pillon, A. (2013). When does action comprehension need motor involvement? Evidence from upper limb aplasia. Cognitive Neuropsychology, 30(4), 253–283. https://doi.org/10.1080/02643294.2013.853655

Vannuscorps, G., Pillon, A., & Andres, M. (2012). Effect of biomechanical constraints in the hand laterality judgment task: where does it come from? Frontiers in Human Neuroscience, 6, 299.

Vannuscorps, G., F Wurm, M., Striem-Amit, E., & Caramazza, A. (2019). Large-Scale Organization of the Hand Action Observation Network in Individuals Born Without Hands. Cerebral Cortex, 29(8), 3434–3444. https://doi.org/10.1093/cercor/bhy212

Verzijl, H. T. F. M., Van der Zwaag, B., Cruysberg, J. R. M., & Padberg, G. W. (2003). Möbius syndrome redefined: A syndrome of rhombencephalic maldevelopment. Neurology, 61(3), 327–333. https://doi.org/10.1212/01.WNL.0000076484.91275.CD

Watkins, K. E., Strafella, A. P., & Paus, T. (2003). Seeing and hearing speech excites the motor system involved in speech production. Neuropsychologia, 41(8), 989–994. https://doi.org/10.1016/S0028-3932(02)00316-0

